# A portable electrochemical DNA sensor for sensitive and tunable detection of piconewton-scale cellular forces

**DOI:** 10.1101/2024.03.24.586508

**Authors:** Mahmoud Amouzadeh Tabrizi, Ahsan Ausaf Ali, Murali Mohana Rao Singuru, Lan Mi, Priyanka Bhattacharyya, Mingxu You

**Author notes:** Corresponding authors E-mail address (M. Amouzadeh Tabrizi), (M. You).

## Abstract

Cell-generated forces are a key player in cell biology, especially during cellular shape formation, migration, cancer development, and immune response. A new type of label-free smartphone-based electrochemical DNA sensor is developed here for cellular force measurement. When cells apply tension forces to the DNA sensors, the rapid rupture of DNA duplexes allows multiple redox reporters to reach the electrode and generate highly sensitive electrochemical signals. The sensitivity of these portable sensors can be further enhanced by incorporating a CRISPR-Cas12a system. Meanwhile, the threshold force values of these DNA-based sensors can be rationally tuned based on the force application geometries and also DNA intercalating agents. Overall, these highly sensitive, portable, cost-efficient, and easy-to-use electrochemical sensors can be powerful tools for detecting different cell-generated molecular forces.

**Graphical Abstract:** 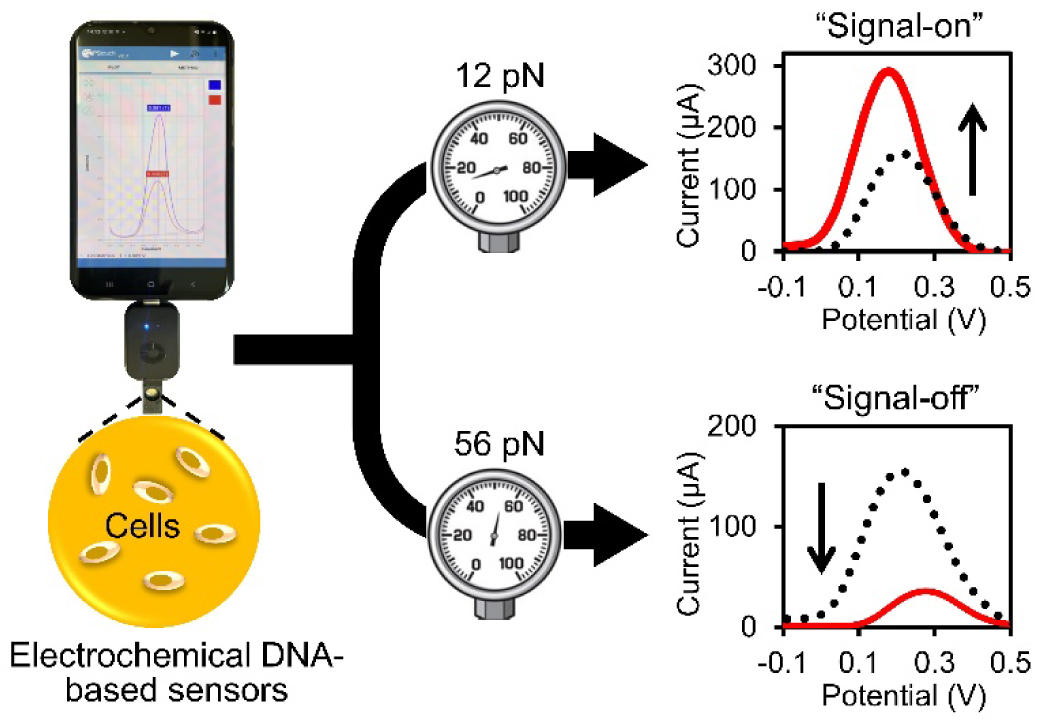

## 1. Introduction

Cells can adhere to the extracellular matrix and other cells during their migration, deformation, invasion, and metastasis processes.^1–3^ These cellular adhesions are often mediated by membrane molecules such as integrins and cadherins,^4–6^ which can not only bind with their target ligands but also generate forces at the piconewton (pN) scale. Thanks to some recent advancements in biotechnologies such as traction force microscopy,^7^ micropillar arrays,^8^ and fluorescent tension probes,^9–12^ our understanding of the biological roles of these cell-generated adhesion forces has been significantly improved during the past years. Among different fluorescent tension probes, various force-sensitive DNAs,^9,10^ polymers,^13,14^ and peptides,^15–17^ have been integrated as the transducers for the detection of molecular level cell-generated adhesion forces. Because DNA-based force probes can be easily synthesized and modified with different ligands and functional moieties, meanwhile, their force threshold values can be modularly and precisely tuned, these fluorescent DNA probes are arguably the most widely used ones for measuring cellular mechanical forces.

We recently developed the first electrochemical DNA-based sensors for the detection of molecular forces generated by live cells.^18^ Compared to fluorescent probes, these new electrochemical force sensors are much more portable, smaller, and simpler to use, which can potentially be adopted by the broader scientific community. However, the sensitivity of our previously designed “first-generation” electrochemical DNA force sensors (Fig. 1a) is relatively low. For example, to detect integrin-mediated adhesion forces, at least ~10^4^ mL^-1^ HeLa cells are still needed. In this current work, we designed a “second-generation” label-free electrochemical DNA-based force sensor with much improved sensitivity and cost-efficiency. Unlike in our previous design where the DNA probe is covalently labeled with a redox reporter, herein, the redox reporters are free in the solution (Fig. 1a). In this kind of “label-free” electrochemical sensors, each force-induced DNA detachment event on the surface of the biosensor will result in multiple redox reporters with decreased mass transfer limitation to reach the electrode and generate electrochemical signals.^19,20^ Thus, compared to “labeled” sensors that exhibit 1:1 force-induced signals, an amplified sensitivity in generating electrochemical force signals can be observed in these new label-free sensors.

**Fig. 1.**
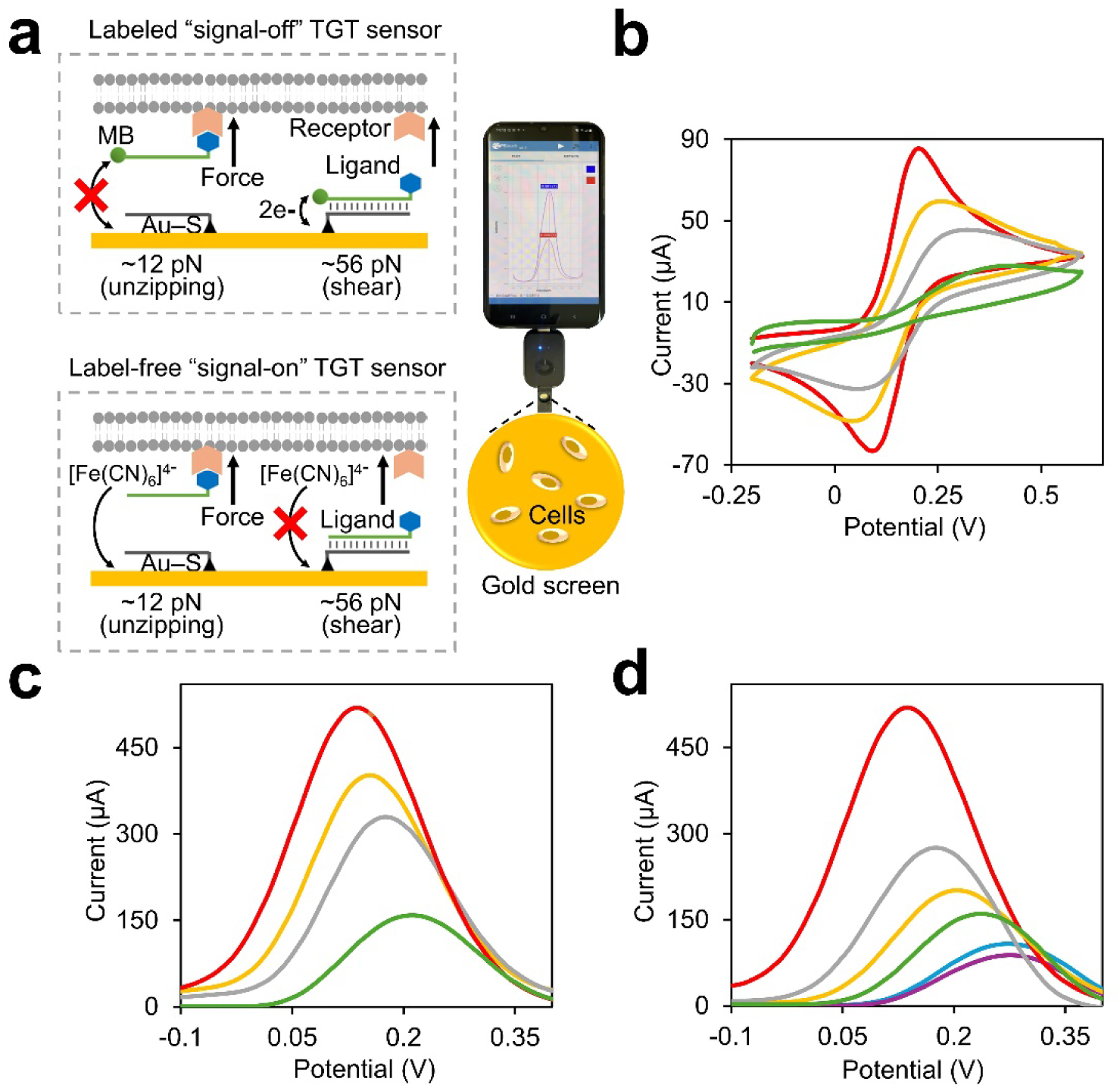
**(a)** Schematic of the smartphone-based labeled or label-free electrochemical tension gauge tether (TGT) force sensors. Cell adhesion forces rupture the double-stranded TGT structures, and as a result, in “signal-off” labeled design, the methylene blue redox reporter exhibited a decreased electrochemical signal, while in “signal-on” label-free system, [Fe(CN)_6_]^4-^ can reach the surface of the electrode to generate an increased signal. **(b)** Cyclic voltammograms and **(c)** square wave voltammograms for characterizing the fabrication process of the rigid TGT sensors. These measurements were performed in a solution containing (v/v) 50% DMEM and 50% phosphate buffer (0.2 M, pH 7.4) and 5.0 mM [Fe(CN)_6_]^4−^, respectively on the unmodified Au-SPE (red line), thiolated anchor DNA-modified Au-SPE (orange line), 1-hexanethiol-passivated thiolated anchor DNA-modified Au-SPE (grey line), and that after adding ligand DNA strand (green line). The cyclic voltammetry signals were recorded at the scan rate of 50 mV·s^−1^. **(d)** Square wave voltammograms for characterizing the fabrication process of the soft TGT sensors as measured in a solution containing (v/v) 50% DMEM and 50% phosphate buffer (0.2 M, pH 7.4) and 5.0 mM [Fe(CN)_6_]^4−^, respectively on the unmodified Au-SPE (red line), 3-mercaptopropionic acid-modified Au-SPE (orange line), after attaching poly-L-lysine (grey line), after adding thiolated anchor DNA (green line), after adding ligand DNA strand (blue line), and that after the BSA blocking (purple line).

In our design, a double-stranded DNA force probe, known as “tension gauge tether” (TGT), is attached to the surface of a gold screen-printed electrode (Au-SPE), which is further incorporated to a portable electrochemical device that is compatible with smartphones. Such a smartphone-based electrochemical sensor can be potentially used as a low-cost point-of-care testing device for the cellular force detection. The TGT probes can be engineered to respond to a wide range (~12–56 pN) of cell-generated molecular forces. Once the cell adhesion rupture the DNA duplex and induce the detachment of the ligand-modifed TGT strand, the diffusibility of the redox reporter, i.e., ferrocyanide [Fe(CN)_6_]^4−^, to the surface of the electrode is significantly increased, causing a positive change in the electrochemical signal. To further improve the sensitivity of the probes, a CRISPR-Cas12a system^21^ is also incorporated to cleave the remaining surface-attached thiolated anchor TGT strand. As a result, the mass transfer limitation of the redox reporter can be additionally decreased to develop a highly sensitive sensor for detecting cell-generated forces. The rupture forces of these TGT sensors can also be simply tuned by adding different amounts of intercalating agents, without changing the sequences of the DNA probes. We expect these powerful electrochemical sensors can be potentially used for measuring various types of cell-generated molecular forces and cell adhesion events.

## 2. Materials and Methods

### 2.1. Reagents and apparatus

Double deionized water (18.6 MΩ·cm^-1^) was used throughout this project. The DNA oligonucleotides were custom synthesized and purified by W. M. Keck Oligonucleotide Synthesis Facility at Yale University School of Medicine, and the sequences are: (1) biotinylated ligand DNA strand: 5’-CACAGCACGGAGGCACGAC AC-biotin-3’; (2) 12 pN anchor DNA strand: 5’-HS-GTGTCGTGCCTCCGTGCTGTG-3’; (3) 56 pN anchor DNA strand: 5’-GTGTCGTGCCTCCGTGCTGTG-SH-3’; (4) crRNA for Cas12a: 5’-CACAGCACGGAGGCACGACAC-3’. The CRISPR-Cas12a (EnGen® Lba Cas12a (Cpf1)) enzyme was brought from New England Biolabs. Hanks’ Balanced Salt Solution (HBSS), 4-(2-hydroxyethyl)-1-piperazine ethane sulfonic acid (HEPES), sodium bicarbonate, potassium chloride (KCl), sodium chloride (NaCl), dipotassium phosphate (K_2_HPO_4_), sulphuric acid (H_2_SO_4_), poly-l-lysine, ethylenediaminetetraacetic acid (EDTA), tris(2-carboxyethyl)phosphine (TCEP), ferrocyanide [Fe(CN)_6_]^4-^, adriamycin, dulbecco’s modified eagle medium (DMEM), streptavidin, biotinylated cyclic arginine-glycine-aspartic acid (B-cRGDfK), sulfosuccinimidyl 4-(N-maleimidomethyl)cyclohexane-1-carboxylate (sulfo-SMCC), 1-hexanethiol, and latrunculin B were obtained from Thermo Fisher Scientific and used without further purification. Gold screen-printed electrodes (Au-SPE) with working electrode made of gold, auxiliary electrode made of platinum, reference electrode and electric contacts made of silver (dimensions: 3.4×1.0×0.05 cm^3^, length×width×height) were purchased from Metrohm-DropSens (Llanera, Spain). The cyclic voltammetry (CV), electrochemical impedance spectroscopy (EIS), and square wave voltammetry (SWV) studies were performed using a Sensit Smart electrochemical device from PalmSens (Houten, Netherlands).

### 2.2. Fabrication of the rigid electrochemical TGT sensor

The fabrication of the electrochemical TGT sensors was similar to that shown in our previous work.^18^ The surface of the Au-SPE was first cleaned using 2 M H_2_SO_4_ solution until no change in the cyclic voltammogram was observed when scanning the potential in the range from −0.3 V to 1.2 V (Fig. S1a). Then, 100 µL of TCEP-reduced 12 pN (or 56 pN) DNA anchor strand (5 µM) was dropped onto the cleaned Au-SPE surface and kept in the refrigerator for 16 h. Following incubation, the DNA-anchored Au-SPE was rinsed with copious amounts of 0.1 M phosphate buffer (pH 7.4) to wash away nonspecifically adsorbed DNA strands. 100 μL of 1-hexanethiol (100 μM) was then dropped on the surface of the electrode to block the remaining active sites on the electrode surface by incubating at 37 °C for 1 h. After that, 100 µL of the biotinylated DNA ligand strand (5.0 µM) was added to the surface of the electrode to generate double-stranded DNA probes by incubating at room temperature for 1 h. The electrode was then washed profusely with 0.1 M phosphate buffer (pH 7.4), and subsequently, 100 µL of 5.0 µM streptavidin was dropped on the surface of electrode to interact with biotinylated DNA duplex at room temperature for 1 h. Again, after rinsing plentifully with 0.1 M phosphate buffer (pH 7.4), 100 µL of 5.0 µM biotinylated cyclic arginine-glycine-aspartic-acid-D-phenylalanine-lysine (cRGDfK) was dropped on the surface of the electrode to interact with streptavidin at room temperature for 1 h. Consequently, the electrode was washed with 0.1 M phosphate buffer (pH 7.4) and 100 µL of 100 µM bovine serum albumin (BSA) was casted on the surface of the electrode to block the remaining active sites. As a final step, the fabricated TGT sensor was rinsed with 0.1 M phosphate buffer (pH 7.4) and stored at 4 °C before usage.

### 2.3. Fabrication of the soft TGT sensor

To fabricate a soft TGT sensor, 100 µL of 5 µM 3-mercaptopropionic acid was first casted on the surface of the cleaned Au-SPE by self-assembly via the Au-S bond. After 16 h of incubation, the electrode was washed with plenty of 0.1 M phosphate buffer (pH 7.4), and then 100 µL of 0.05 M PBS of pH 7.0 containing 10 mM 1-ethyl-3-(3-dimethylaminopropyl)carbodiimide (EDC) and 20 mM N-hydroxysuccinimide (NHS) was dropped on the surface to activate the carboxylic acid group of the 3-mercaptopropionic acid for 1 h. Consequently, the electrode was washed with phosphate buffer (pH 7.4) and 100 µL of 0.1 mg/mL poly-L-lysine was casted on the surface to attach via forming amide groups. After 2 h incubation, the electrode was again washed with phosphate buffer (pH 7.4), and then 100 µL of 0.5 mM sulfo-SMCC was casted on the surface for 2 h to attach to the amine groups of poly-L-lysine.^22^ After washing, 100 µL of 5 µM thiolated anchor DNA strand was dropped on the surface of the electrode to conjugate with the primary amine groups of the poly-L-lysine. Afterwards, additional washing was performed with 0.1 M PBS (pH 7.4) and then 100 µL of 5.0 µM biotinylated ligand DNA strand was added to generate double-stranded DNA on the surface of the electrode at room temperature for 1 h. After washing profusely with 0.1 M phosphate buffer (pH 7.4), 100 µL of 5.0 µM streptavidin was added at room temperature for 1 h. Again, the electrode was rinsed plentifully with 0.1 M phosphate buffer (pH 7.4), and then 100 µL of 5.0 µM biotinylated cyclic arginine-glycine-aspartic acid (B-cRGDfK) was dropped on the surface for a 1 h incubation at room temperature. As a final step, 100 µL of 1% BSA solution in 0.1 M phosphate buffer (pH 7.4) was casted on the surface of the electrode to avoid any non-specific binding during the cellular measurements. Such fabricated soft TGT sensors were rinsed with 0.1 M phosphate buffer (pH 7.4) and stored at 4 °C before usage.

### 2.4. Measurement of cell-generated forces

HeLa cells were cultured in DMEM with 10% fetal bovine serum, 100 U/mL penicillin, and 100 U/mL streptomycin in an Eppendorf Galaxy incubator at 5% (v/v) CO_2_. Before measurement, the cells were first detached by adding 2 mM EDTA solution, consisting of 1×HBSS, 0.06% sodium bicarbonate, and 0.01 M HEPES (pH 7.6), for 10 min. The solution was then centrifuged three times at 1,200 rpm for 7 min and re-suspended in a measuring solution containing (v/v) 50% DMEM and 50% phosphate buffer (0.2 M, pH 7.4), with a final concentration from ~100 cells/mL to 1×10^6^ cells/mL. In a typical measurement, ~1×10^5^ HeLa cells (100 µL) were added on the surface of the above-prepared TGT electrochemical sensor and allowed the cells to interact with the DNA probes for 75 min. After that, the electrode was washed with 0.1 M phosphate buffer (pH 7.4) and added with 100 µL of 5 mM [Fe(CN)_6_]^4−^ on the surface to start recording the SWV signals of the [Fe(CN)_6_]^4−^. SWV parameters were as follows: step potential, 20 mV; pulse amplitude, 50 mV; and frequency, 20 Hz. The impact of different experimental conditions on the response of the TGT sensors were also investigated as shown below.

### 2.5. The treatment of cells with a force inhibition drug

We have studied how the latrunculin B treatment can influence the cellular force generation. For this purpose, ~1×10^6^/mL HeLa cells were first pretreated with 5–60 µM of latrunculin B for 60 min at 37 °C inside a cell culture incubator. After that, cells were separated and dispersed in cell culture media, ~1×10^5^ cells was then casted onto the surface of the TGT sensor and incubated for 75 min. After washing with 0.1 M phosphate buffer (pH 7.4) for several times, the SWV of 5 mM [Fe(CN)_6_]^4−^ on the electrode was recorded using the above-mentioned parameters: step potential, 20 mV; pulse amplitude, 50 mV; and frequency, 20 Hz.

### 2.6. cRGD treatment of the cells

To study the specificity of the sensor in detecting integrin-mediated force generation, we have added 1.0, 2.5, or 5.0 µM of free cyclic arginine-glycine-aspartic acid (cRGD) molecules with ~1×10^6^/mL HeLa cells and allowed the RGD to attach with the cell membrane integrins for 60 min. As a result, these pre-occupied integrins of the HeLa cells cannot interact with the TGT sensors any more. After the treatment, the cells were separated and dispersed in the cell culture media, ~1×10^5^ cells were then added to the surface of the TGT sensor and allowed to interact for 75 min. Finally, the electrode was washed with 0.1 M phosphate buffer (pH 7.4) for several times and the SWVs of 5 mM [Fe(CN)_6_]^4−^ were recorded under the following condition: step potential, 20 mV; pulse amplitude, 50 mV; and frequency, 20 Hz.

### 2.7. Intercalation of doxorubicin in the TGT sensor

Modified based on a previously reported protocol,^23^ 100 µL of 5 mg/mL doxorubicin was casted on the surface of the 12 pN TGT sensor and allowed to intercalate inside the DNA duplex at 37 °C for 30 min. After that, the electrode was washed with 0.1 M phosphate buffer (pH 7.4) for several times and stored at 4 °C before usage. The recorded cyclic voltammograms at different scan rates indicated that doxorubicin was capable of intercalating inside the TGT probes, as the peak current values exhibited a linear relationship with the scan rates (Fig. S2).^24^ To determine the number of intercalated doxorubicin in each DNA duplex, we first calculated the surface coverage of double-stranded DNA (Γ_dsDNA_) by integrating the reduction peak of [Ru(NH_3_)_6_]^3+^, which interacted with the phosphate groups of the DNA. As shown in Table S1 and Fig. S3, the surface coverage of [Ru(NH_3_)_6_]^3+^ was found to be ~3.5×10^-10^ mol/cm^2^. Considering the charge of [Ru(NH_3_)_6_]^3+^ (z = 3) and the number of nucleotides (m = 42), Γ_dsDNA_ was calculated based on Γ_ds DNA_ = Γ_Ru_×(z/m) as ~2.5×10^-11^ mol/cm^2^. The surface coverage of doxorubicin (Γ_DOX_) was similarly calculated to be ~8.0×10^-11^ mol/cm^2^ by integrating the reduction peak of doxorubicin using the CV method. By comparing the Γ_dsDNA_ and Γ_DOX_, ~3.2 doxorubicin were found to be inserted into each TGT probe.

### 2.8. Cas12a-mediated signal amplification of the TGT sensors

After rupturing the ligand DNA strand of the TGT sensors by the cells, the Cas12a/crRNA conjugate was used to further cleave the remaining anchor DNA strand to additionally improve the diffusibility of [Fe(CN)_6_]^4−^ and to increase the sensitivity of the sensor. Here, the Cas12a/crRNA conjugate was prepared based on the method reported previously in the literature.^25,26^ Briefly, 100 nM CRISPR-Cas12a, 100 nM crRNA, and 1×NEBufer 2.1 were first mixed for 30 min. After that, 100 µL of the mixture at 30, 60, 90, or 120 nM concentration was casted on the surface of the cell-incubated electrode or just anchor DNA strand-conjugated Au-SPE for 30 min. During this process, the anchor DNA was cleaved. After washing the electrode with 0.1 M phosphate buffer (pH 7.4) for several times, the SWVs of 5 mM [Fe(CN)_6_]^4−^ were recorded using the following parameters: step potential, 20 mV; pulse amplitude, 50 mV; and frequency, 20 Hz.

### 2.9. Rupture force estimation of the TGT sensors

The tension tolerance (T_tol_) of the TGT probes depends on the orientation of the applied forces and the length of the duplex region. T_tol_ is defined as the rupture force required to unfold 50% of the DNA duplex. Using the de Gennes model,^27,28^ T_tol_ is calculated as T_tol_= 2F_c_·[X^-1^·tanh(X·L/2) + 1]. Here, F_c_ is the rupture force of each base pair (3.9 pN), L is the base pair number between two force-applying points on the complementary TGT strands, and X = (2R/Q)^1/2^, measuring the DNA duplex elasticity based on two spring constants between neighbors in a strand (Q) or between base pairs in a duplex (R). Under conditions where a constant force is applied for 1–2 s, X^−1^= 6.8.

## 3. Results and Discussion

### 3.1. Design and fabrication of the TGT sensors

Using previously reported DNA duplex sequences,^18,29^ we first anchored a TGT force probe with T_tol_ of 12 pN onto the Au-SPE surface. The whole fabrication process was characterized in each step using cyclic voltammetry (CV) and square wave voltammetry (SWV) in a measuring solution containing (v/v) 50% DMEM and 50% phosphate buffer (0.2 M, pH 7.4) and 5.0 mM [Fe(CN)_6_]^4−^ redox reporter. As shown in Fig. 1b and S1, the CV curves of [Fe(CN)_6_]^4−^ on the Au-SPE electrode were measured at a scan rate of 50 mV·s^-1^. After initial assembly of the thiolated anchor DNA strand to the surface of the Au-SPE, the peak current intensities of [Fe(CN)_6_]^4−^ decreased from I_pa_ ~ 85 μA and I_pc_ ~ −60 μA to I_pa_ ~ 67 μA and I_pc_ ~ −52 μA. Meanwhile, the peak-to-peak separation (ΔE) increased from ~0.13 V to ~0.16 V, indicating an enhanced mass transfer limitation of [Fe(CN)_6_]^4−^ at the surface of Au-SPE due to the electrostatic repulsion between the negatively charged DNA and negatively charged [Fe(CN)_6_]^4−^ redox reporter. After further blocking the remaining active sites of the Au-SPE surface with 1-hexanethiol and then added the complementary ligand DNA strands, the mass transfer limitation of [Fe(CN)_6_]^4−^ increased more, and consequently, its peak intensities decreased to I_pa_ ~ 49 μA, I_pc_ ~ −31 μA and I_pa_ ~ 28 μA, I_pc_ ~ −6 μA, respectively, and the ΔE values increased to ~0.19 V and ~0.41 V (Fig. 1b).

The interface properties of the electrodes were also measured with the SWV method. After the coating of DNA and 1-hexanethiol onto the surface of Au-SPE, due to the mass transfer limitation for [Fe(CN)_6_]^4−^, the peak current intensity of the signals (ΔI = I_peak_ – I_background_) decreased step by step from ~480 µA to ~150 µA (Fig. 1c). These SWV results are coherent with the CV data.

Moreover, as the softness of the surface can also affect the force response of the TGT sensors,^30,31^ we have also fabricated a type of soft TGT sensors and characterized using SWV. As shown in Fig. 1d, the intensity of the peak current first decreased from ~480 µA to ~200 µA after adding 5 µM of 3-mercaptopropionic acid to self-assemble on the surface of the Au-SPE. While after attaching poly-L-lysine, the peak current intensity increased to ~275 µA as the positively charged amine groups within the poly-L-lysine could adsorb the negatively charged [Fe(CN)_6_]^4−^. After further adding the anchor DNA strand and ligand DNA strand, the intensity of the peak current again decreased to ~160 µA and ~115 µA, due to the electrostatic repulsion between DNA and [Fe(CN)_6_]^4−^. Finally, after blocking the surface with 100 µM BSA, the intensity of the signal decreased to <90 µA, indicating the successful fabrication of soft TGT sensors on the electrode.

### 3.2. TGT-based detection of cell adhesion forces

We next applied such fabricated TGT sensors to detect cell-generated adhesion forces. By modifying the cRGDfK ligand on the 12 pN TGT probes, integrin αvβ3-mediated surface attachement of HeLa cells were studied here. After incubating ~1×10^5^ HeLa cells on the surface of either soft or more rigid electrode for 90 min, the peak current intensity of the TGT sensors clearly increased in both cases (Fig. 2a, 2b, and S4a), which is expectedly due to the rupture of ligand DNA strands by the HeLa cells that results in more [Fe(CN)_6_]^4−^ redox reporters reaching the surface of the electrode. Our results indicated that the cellular response of soft TGT sensor is much lower compared to the rigid electrode (Fig. 2c). These more rigid TGT sensors will be used throughout this work. As the SWV signals barely changed after 75 min, for the following studies, 75 min will be applied as the optimum incubation time between the TGT sensor and HeLa cells.

**Fig. 2.**
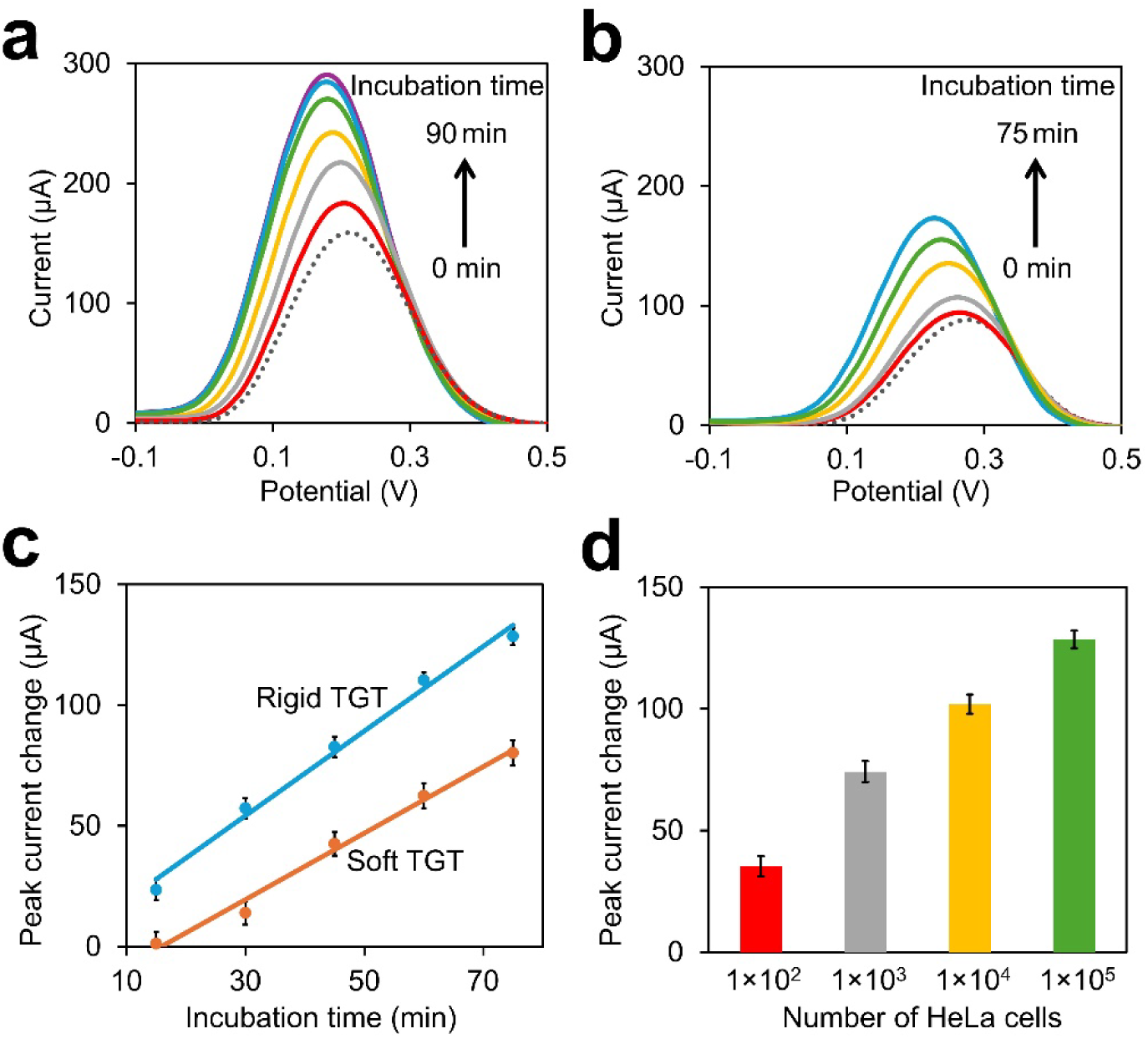
**(a)** Square wave voltammetries of the rigid TGT sensor before (dotted line) and after adding ~1×10^5^ HeLa cells for 15, 30, 45, 60, 75, and 90 min, respectively. **(b)** Square wave voltammetries of the soft TGT sensor before (dotted line) and after adding ~1×10^5^ HeLa cells for 15, 30, 45, 60, and 75 min, respectively. **(c)** Cell incubation time-dependent changes in the peak current values as measured using the rigid or soft TGT sensors. **(d)** Peak current changes of the rigid TGT sensor after adding from ~100 to ~1×10^5^ HeLa cells for 75 min. All these measurements were performed in a solution containing (v/v) 50% DMEM and 50% phosphate buffer (0.2 M, pH 7.4) and 5.0 mM [Fe(CN)_6_]^4−^. The step potential was set as 20 mV, the pulse amplitude was at 50 mV, and the frequency was at 20 Hz. Shown are the mean and standard error peak values after subtracting the background signals from four replicated tests.

The effect of HeLa cell concentrations on the TGT electrochemical signals were studied next. As can be seen in Fig. 2d and S4b, after incubating 100 to 1×10^5^ HeLa cells on the TGT sensors for 75 min, the SWV signals obviously increased as higher concentrations of the cells were added to the electrode, indicating more ligand DNA strands were ruptured by the cells. We further compared these results with our previous design using methylene blue-labeled 12 pN TGT sensors.^18^ The sensitivity of these new label-free DNA sensors was ~140 times higher than that of the labeled ones (Fig. S5). It is worth noting that the current label-free TGT sensors can provide a type of “signal-on” response, while the previous labeled sensors are based on a “signal-off” mechanism.

To validate whether the integrin-RGD interactions are responsible for the observed TGT electrochemical signals, we blocked the integrins on the HeLa cell membranes by first adding 1–5 µM of free cyclic arginine-glycine-aspartic acid (cRGD) molecules (Fig. 3a and S6a). Indeed, compared to the cell adhesion signals without the cRGD treatment, much decreased peak current intensities were observed. After the treatment with 5 µM cRGD, the SWV signals of the TGT sensors were almost identical to that without adding the cells.

**Fig. 3.**
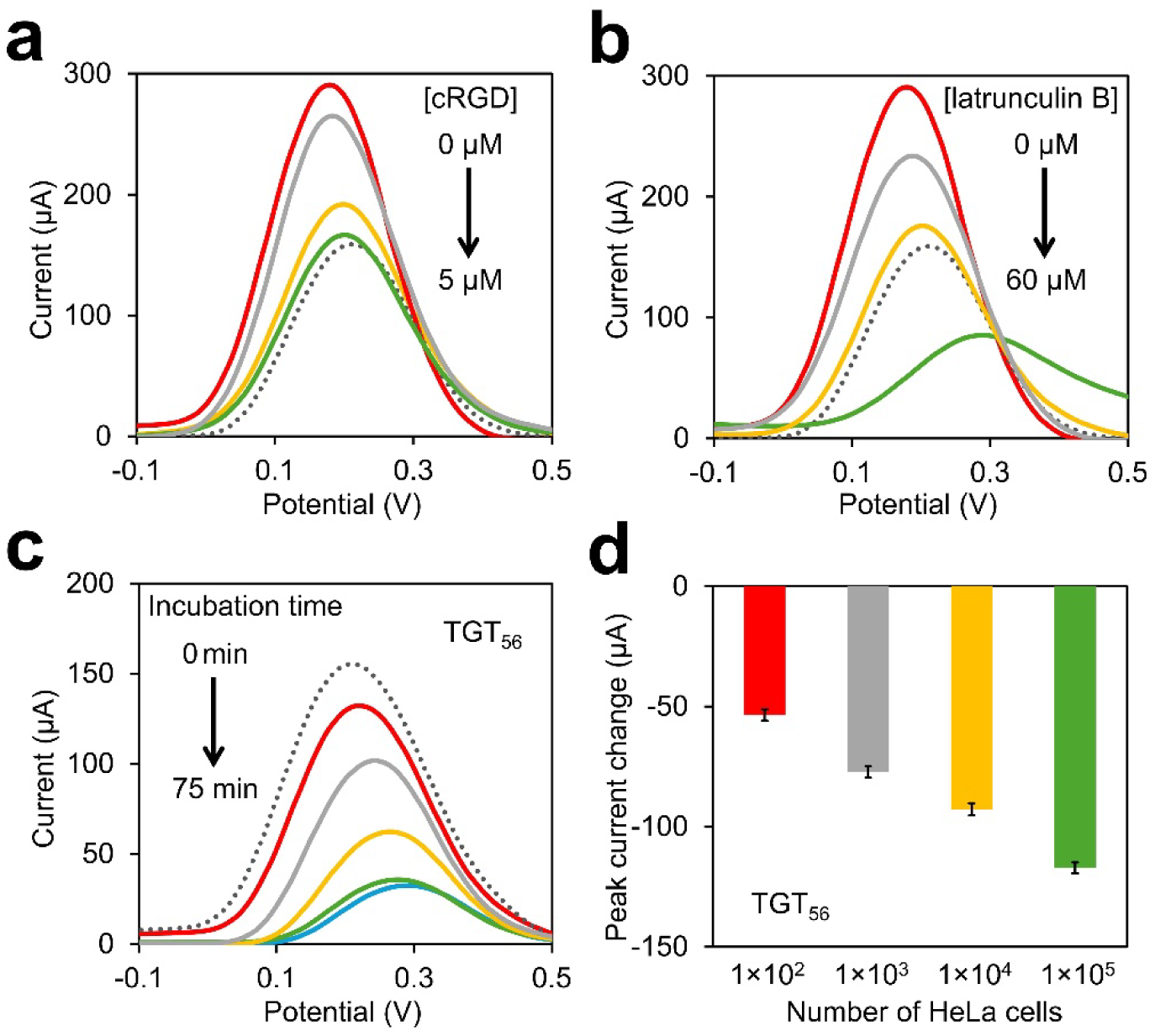
**(a)** Square wave voltammetries of the 12 pN TGT sensor in the absence of HeLa cells (dotted line) or after adding ~1×10^5^ HeLa cells for 75 min. These cells have been pre-treated with 0, 1, 2.5, or 5 µM of cyclic arginine-glycine-aspartic acid (cRGD) for 75 min. **(b)** Square wave voltammetries of the 12 pN TGT sensor in the absence of HeLa cells (dotted line) or after adding ~1×10^5^ HeLa cells for 75 min. These cells have been pre-treated with 0, 5, 30, or 60 µM latrunculin B for 60 min. **(c)** Square wave voltammetries of the 56 pN TGT sensors before (dotted line) and after adding ~1×10^5^ HeLa cells for 15, 30, 45, and 60 min, respectively. **(d)** Peak current changes of the 56 pN TGT sensors after adding from ~100 to ~1×10^5^ HeLa cells for 60 min. All these measurements were performed in a solution containing (v/v) 50% DMEM and 50% phosphate buffer (0.2 M, pH 7.4) and 5.0 mM [Fe(CN)_6_]^4−^. The step potential was set as 20 mV, the pulse amplitude was at 50 mV, and the frequency was at 20 Hz. Shown are the mean and standard error peak values after subtracting the background signals from four replicated tests.

To further test if the electrochemical TGT signals are indeed due to the cell-generated forces, we treated HeLa cells with 5–60 µM of latrunculin B, a force-inhibiting drug that prevents actin polymerization and also the transition of cellular G-actins into F-actins.^32^ As shown in Fig. 3b and S6b, when ~1×10^5^ HeLa cells were pre-treated with 5 µM or 30 µM of latrunculin B, much less DNA probes were ruptured from the surface of the electrode, resulting in a significantly decreased peak current intensity from ~290 µA to ~180 µA. All these data suggested that the TGT electrochemical signals are resulted from the integrin-RGD interactions, which can generate forces to rupture the DNA duplex and change the mass transfer limitation of the redox reporter for the detection of cell adhesion forces.

### 3.3. Detecting cellular adhesion events using the TGT sensors

While interestingly, in the case of 60 µM latrunculin B treatment, the SWV signals were even lower than that in the absence of the cells. The reasonable explanation can be that under such a strong force inhibition condition, HeLa cells will just attach to the surface of the electrode via the integrin-RGD interactions, rather than force-mediated rupturing of the RGD-modified ligand DNA strands. As a result, the SWV signals of [Fe(CN)_6_]^4−^ were decreased to a greater extent due to mass transfer limitation caused by electrode surface-attached HeLa cells.

We further studied this phenomena by designing another TGT sensor that will be ruptured by forces ≥56 pN, termed “TGT_56_”. TGT_56_ shared the same sequence with the above-used 12 pN TGT sensor, but the thiolated anchor group is located at the opposite end of the ligand moiety. Such a “shear mode” design is known to increase the force threshold value of the TGT sensors,^29^ and our previous results suggested that integrin-generated forces during the HeLa cell attachement is not large enough to rupture these 56 pN TGT structures.^18^ As shown in Fig. 3c and S7a, after adding ~1×10^5^ HeLa cells on the surface of the TGT_56_ sensor, the peak current intensity continuously decreased during the first 60 min incubation time.

Moreover, we investigated the impacts of HeLa cell number (from ~100 to 1×10^5^ cells) on the TGT_56_ signals. Our results indicated that as the concentration of the HeLa cells getting increased, the SWV signals of the sensors kept decreasing (Fig. 3d and S7b). These label-free TGT_56_ sensors provide a “signal-off” response for detecting different concentrations of cancer cells. Unlike 12 pN TGT sensors, the HeLa cells could not rapture the ligand DNA strands of TGT_56_ and thus instead, these cells will attach to the electrode surface through strong binding between integrins and RGDs. Hence, the mass transfer limitation of [Fe(CN)_6_]^4−^ to the surface dramatically increased and resulted in a decreased electrochemical signal, similar to that shown above in the case of 60 µM latrunculin B treatment (Fig. 3b and S6b). All these results collectively demonstrated that by tuning the threshold value of the TGT sensors, these electrochemical devices can be used to detect cell-generated forces and/or cellular adhesion events.

### 3.4. CRISPR-Cas12a-incorporated TGT force sensors

To further improve the sensitivity of these TGT sensors, we have incorporated a CRISPR-Cas12a (Cpf1)/crRNA ribonucleoprotein complex to additionally amplify the electrochemical signals of the DNA sensors. We reason that in our above-designed TGT sensors, even after the cellular rupture and removal of the anchor DNA strands, the remaining anchor DNAs on the surface of the electrode can still repulse some [Fe(CN)_6_]^4−^ redox reporters and prevent them from reaching the electrode to generate electrochemical signals. Inspired by some recently developed Cas12a/crRNA sensors,^12,25,26^ in our new system, as shown in Fig. 4a, by inserting an activator sequence in the anchor DNA strand, in the presence of cellular forces, the rupture of DNA duplex triggers the Cas12a/crRNA complex to recognize the activator sequence and then cleave the surface-attached anchor DNA strands on the electrode. As a result, the electrostatic repulsion between [Fe(CN)_6_]^4−^ and the electrode surface is reduced, which consequently increased the sensitivity of the TGT sensor in detecting molecular tension forces.

**Fig. 4.**
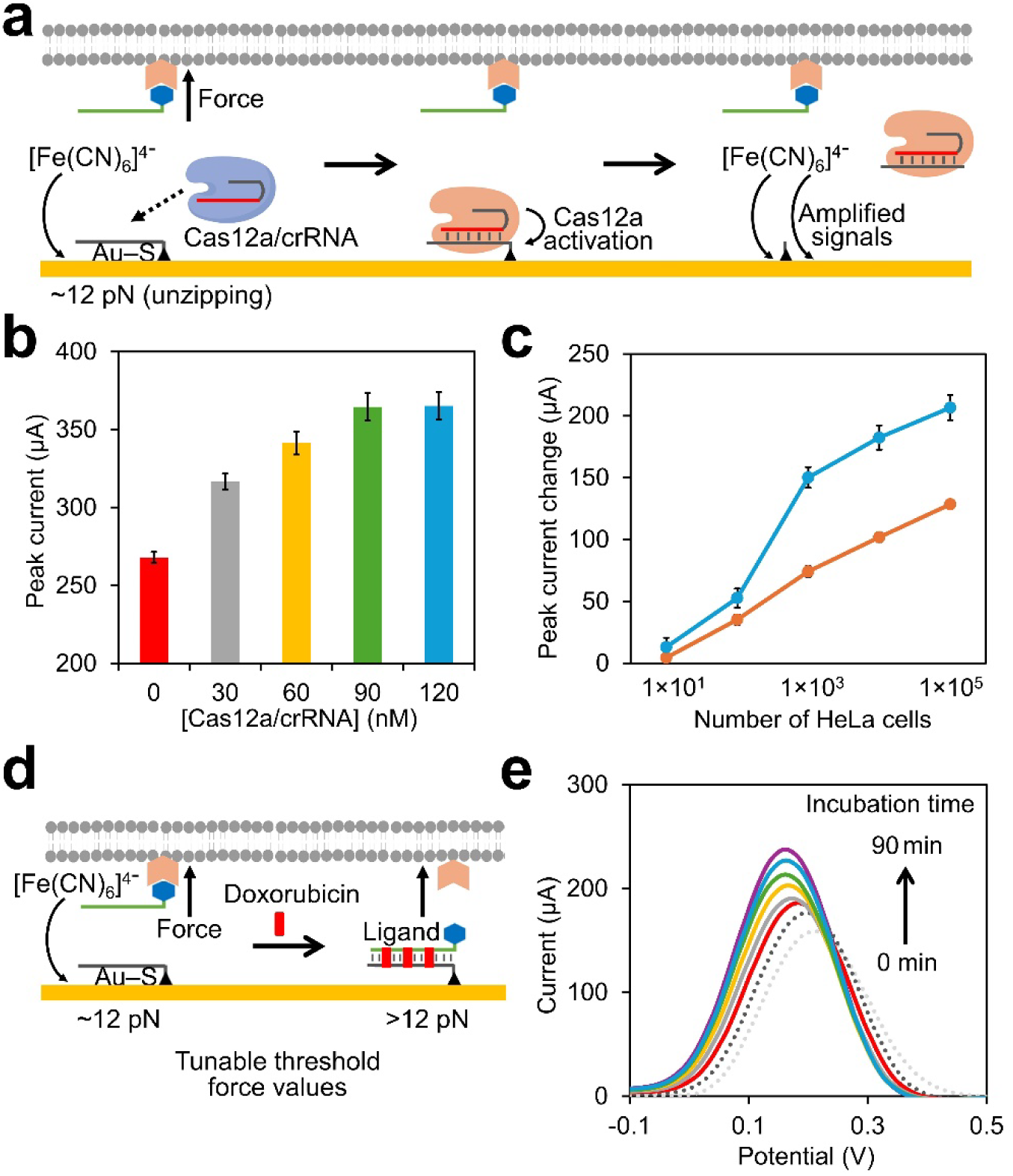
**(a)** Schematic of the Cas12a/crRNA-mediated signal amplification for the TGT force sensors. Upon experiencing cellular forces, the rupture of DNA duplex binds and activates the Cas12a/crRNA complex to cleave the surface-attached anchor DNA strands on the electrode. More [Fe(CN)_6_]^4−^ can reach the surface of the electrode to generate electrochemical signals. **(b)** Peak current of the 12 pN TGT sensors after first incubating with ~1×10^5^ HeLa cells for 75 min, and then adding different concentrations of the Cas12a/crRNA complex for 30 min. **(c)** Peak current change of the 12 pN TGT sensor after adding from ~10 to ~1×10^5^ HeLa cells for 75 min, in the presence (blue line) or absence (orange line) of 90 nM Cas12a/crRNA complex. **(d)** Schematic of the doxorubicin-intercalated TGT sensors with tunable rupture force threshold values. **(e)** Square wave voltammetries of the doxorubicin-intercalated TGT sensor before (black dotted line) and after adding ~1×10^5^ HeLa cells for 15, 30, 45, 60, 75, and 90 min, respectively. As a comparison, light grey dotted line indicates the 12 pN TGT signals without doxorubicin before adding the HeLa cels. All these measurements were performed in a solution containing (v/v) 50% DMEM and 50% phosphate buffer (0.2 M, pH 7.4) and 5.0 mM [Fe(CN)_6_]^4−^. The step potential was set as 20 mV, the pulse amplitude was at 50 mV, and the frequency was at 20 Hz. Shown are the mean and standard error peak values after subtracting the background signals from four replicated tests.

We first studied the effects of Cas12a/crRNA complex concentrations on the SWV signals of [Fe(CN)_6_]^4−^. As can be seen in Fig. 4b and S8a, after incubating ~1×10^5^ HeLa cells on the 12 pN TGT sensors for 75 min, further increased peak current intensities of the sensors were indeed observed as increasing concentrations of the Cas12a/crRNA complex were added from 0 to 90 nM. In the presence of 90 nM Cas12a/crRNA, we further measured the forces generated by different numbers (from ~10 to 1×10^5^) of the HeLa cells (Fig. 4c and S8b). Our results indicated that these CRISPR-Cas12a-powered label-free TGT sensors are ~65% more sensitive than that without adding the Cas12a/crRNA complex, i.e., ~230 times higher sensitivity as compared to our previously developed methylene blue-labeled TGT sensors. Adhesion forces generated by ≤10 cells could now be detected with high precision and accuracy (Fig. 4c).

### 3.5. Tunable rupture forces of the TGT sensors

The force threshold values of the TGT sensors can be adjusted by changing the relative position between the anchor group and ligand moiety, as shown in above-demonstrated 12 pN and 56 pN TGT sensors. However, once these DNA strands were synthesized and immobilized on the surface of the electrode, their force threshold cannot be easily modified. Herein, we wondered for a given “unzipping mode” 12 pN TGT sensor, by simply adding intercalating agents, such as doxorubicin, whether their force-responding signals can be rationally tuned.

Doxorubicin is known to intercalate between the nearby G–C base pairs in double-stranded DNA and increase the rapture forces.^23^ Since there are four (G–C)_2_ base pairs in our 12 pN TGT sequence, up to four doxorubicin molecules may be intercalated in each DNA probe. In our case, we casted 5 mg/mL doxorubicin on the surface of the TGT sensors at 37 °C for 30 min. Under this condition, >3.2 doxorubicin was found to be intercalated inside each double-stranded DNA duplex on the electrode surface (Fig. 4d, S3 and Table S1). After adding ~1×10^5^ HeLa cells onto these doxorubicin-containing TGT sensors and incubated for 90 min, the force-induced [Fe(CN)_6_]^4−^ signal enhancement can sill be clearly observed, however, as compared to that without the doxorubicin treatment, a much slower (>2-fold) increase in the peak current intensities were shown (Fig. 4e). This result can be expected as considering the strong doxorubicin interactions with nearby G–C base pairs, it now becomes more difficult for the HeLa cells to rupture the DNA duplexes. The rupture forces of the TGT sensors can be directly tuned by adding different amounts of the intercalating agents.

## 4. Conclusion

In summary, a type of label-free electrochemical DNA-based force sensor has been fabricated in this project to detect cell-generated molecular tension forces. Using integrin-RGD interaction-mediated HeLa cell adhesion as an example, our results indicated that these novel electrochemical sensors could be used for highly sensitive, versatile and low-cost investigation of mechanicals forces and cellular adhesion events. By changing the distance and relative location between the anchor site and ligand moiety, or by simply adding DNA intercalating agents, these modular sensors can measure forces at tunable threshold values. Moreover, these rapid and easy-to-use smartphone-based electrochemical devices can be straightforwardly applied to study the impacts of different cell incubation time, cell concentrations, and force inhibiting drugs on the cellular force levels. The electrochemical signals and sensitivities of these DNA-based sensors could further be amplified using a CRISPR-Cas12a system. We expect a potential broad usage of these sensitive, portable, and robust tools in helping researchers to study cellular mechanosensing and mechanotransduction.

## Acknowledgments

The authors gratefully acknowledge the support from NIH R35GM133507, Camille Dreyfus Teacher-Scholar Award, SLAS Graduate Education Fellowship, and Paul Hatheway Terry Scholarship. The authors also thank other members of the You Lab for useful discussion and valuable comments.

## Supporting Information Available

Supplementary data associated with this article can be found in the online version.

