## Supplementary Information for "A portable electrochemical DNA sensor for sensitive and tunable detection of piconewton-scale cellular forces"

* Corresponding authors

### Supplementary Figures

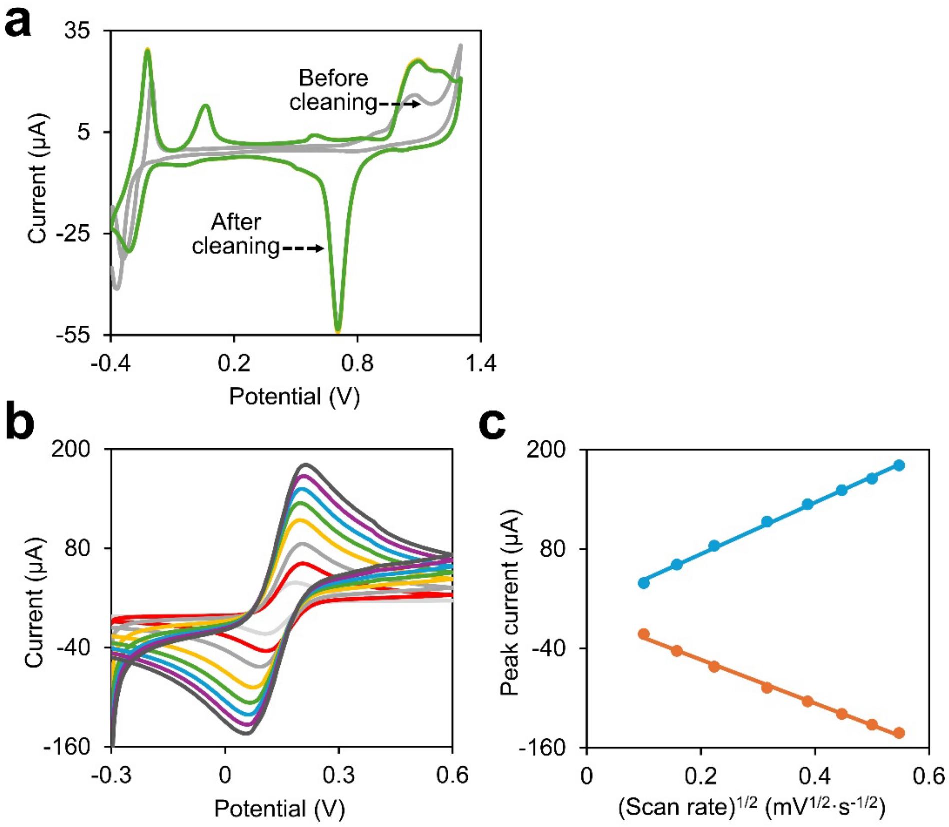


**Fig. S1. (a)** Cyclic voltammogram of the Au-SPE before (light grey line) and after cleaning with 2.0 M $H_2SO_4$ solution, the green and orange lines are obtained from the last two scans. The scan rate was at 10.0 $mV\cdot s^{-1}$ in a 0.1 M phosphate buffer (pH 7.4). **(b)** Cyclic voltammogram of $[Fe(CN)_6]^{4-}$ in a 0.1 M phosphate buffer (pH 7.4) at various scan rates of 10, 25, 50, 100, 150, 200, 250, and 300 $mV\cdot s^{-1}$ from inner to outer. **(c)** A linear correlation was observed between both anodic (orange line) and cathodic (blue line) peak currents ($I_p$) and the square root of scan rates (ν).

Based on the data shown in Fig. S1b, the slope of the anodic log($I_{pa}$) versus log(v) for the Au-SPE was ~0.45 $\mu A/(mV\cdot s^{-1})$, which is quite close to the theoretical value of 0.5 in the case of an ideal diffusion-controlled reaction.[1] These results indicated that the electrochemical kinetics are controlled by the diffusion of $[Fe(CN)_6]^{4-}$ to the surface of the Au-SPE. By drawing the $I_p$ versus the square root of scan rate ($\upsilon^{1/2}$), as shown in Fig. S1c, the electro-active area of the electrode ($A_{eas}$) was also obtained in this case by using the Randles–Sevcik equation for quasi-reversible electrochemical processes:[2] $I_p^{quasi} = \pm 0.436\times F\times A_{eas}\times C\times\sqrt{\frac{n\times F\times D\times \upsilon}{R\times T}}$. Here, $I_p^{quasi}$ is the peak current value, n is the number of electrons (n=1), F is the Faraday constant (96,485 $C\cdot mol^{-1}$), C is the concentration of $[Fe(CN)_6]^{4-}$ ($5.0\times 10^{-6}$ $mol\cdot cm^{-3}$), D is the diffusion coefficient ($7.6\times 10^{-6}$ $cm^2\cdot s^{-1}$), υ is the scan rate (10–300 $mV\cdot s^{-1}$), R is the ideal gas constant (8.314 $J\cdot K^{-1}\cdot mol^{-1}$), and T is the temperature (298 K). Here, the $A_{eas}$ values for the Au-SPE was found to be ~0.085 $cm^{-2}$.

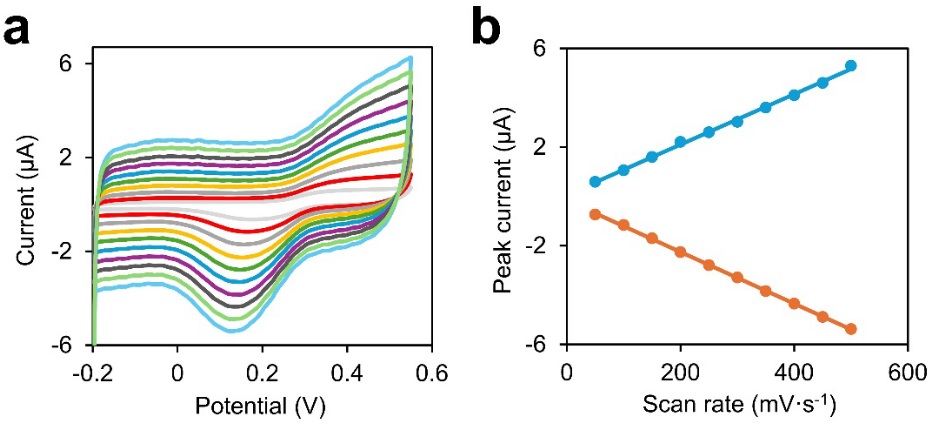


**Fig. S2.** **(a)** Cyclic voltammogram of the intercalated doxorubicin in the 12 pN TGT sensors in a 0.1 M phosphate buffer (pH 7.4) at various scan rates of 50, 100, 150, 200, 250, 300, 350, 400, 450, and 500 mV·s$^{-1}$ from inner to outer. **(b)** A linear correlation was observed between both anodic (orange line) and cathodic (blue line) peak currents ($I_p$) and the scan rates (ν).

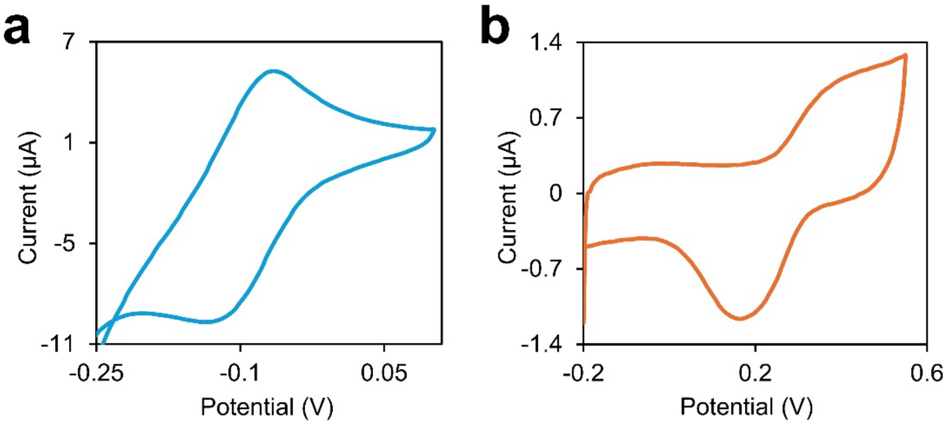


**Fig. S3.** **(a)** Cyclic voltammogram of $[Ru(NH_3)_6]^{3+}$ for determining the surface coverage of double-stranded DNA. Scan was performed at a rate of 100 mV·s$^{-1}$ in a 0.1 M phosphate buffer (pH 7.4). **(b)** Cyclic voltammogram of the intercalated doxorubicin in the 12 pN TGT sensors in a 0.1 M phosphate buffer (pH 7.4) at a rate of 100 mV·s$^{-1}$. From these CV curves, we can obtain the reduction peak current, including non-Faradic current and Faradic current, of both the attached $[Ru(NH_3)_6]^{3+}$ and the intercalated doxorubicin to the 12 pN TGT sensors.

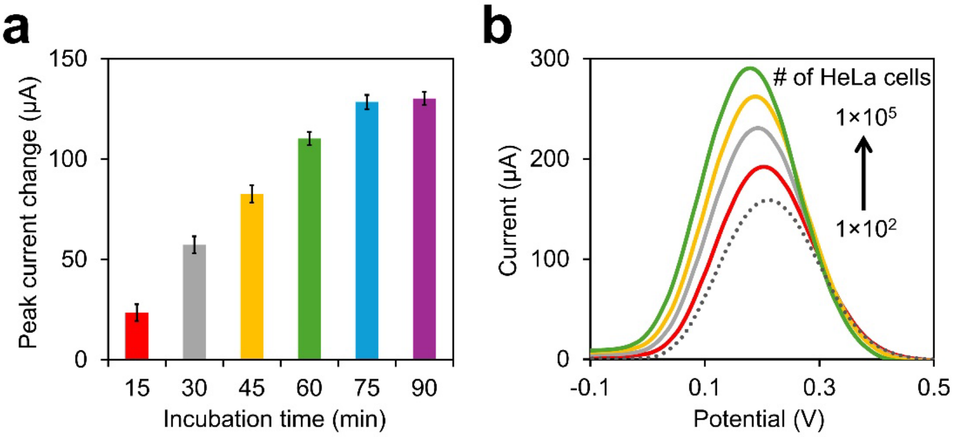


**Fig. S4.** **(a)** Cell incubation time-dependent changes in the peak current values as measured using the rigid 12 pN TGT sensors. **(b)** Square wave voltammetries of the rigid 12 pN TGT sensor before (dotted line) and after adding ~100, ~$1\times10^3$, ~$1\times10^4$, or ~$1\times10^5$ HeLa cells for 75 min. All these measurements were performed in a solution containing (v/v) 50% DMEM and 50% phosphate buffer (0.2 M, pH 7.4) and 5.0 mM $[Fe(CN)_6]^{4-}$. The step potential was set as 20 mV, the pulse amplitude was at 50 mV, and the frequency was at 20 Hz. Shown are the mean and standard error peak values after subtracting the background signals from four replicated tests.

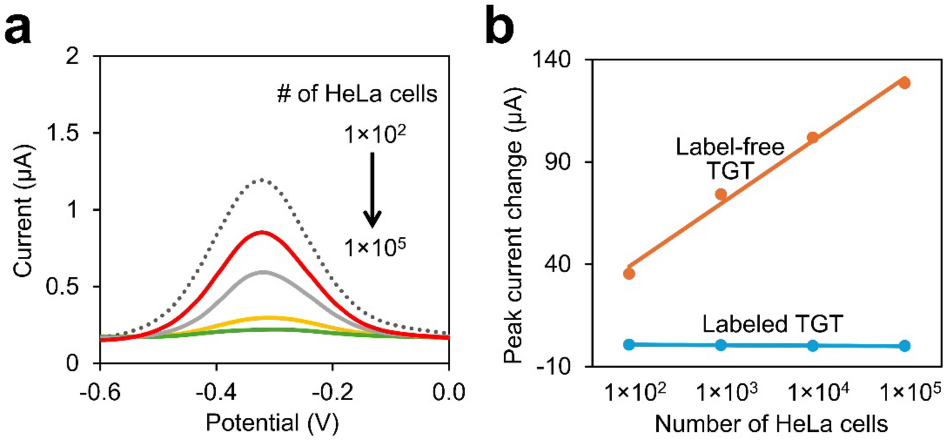


**Fig. S5.** **(a)** Square wave voltammetries of the methylene blue-labeled 12 pN TGT sensor before (black dotted line) or after adding ~100, ~$1\times10^3$, ~$1\times10^4$, or ~$1\times10^5$ HeLa cells for 75 min. **(b)** Peak current changes of labeled and label-free 12 pN TGT sensors after adding from ~100 to ~$1\times10^5$ HeLa cells for 75 min. All these measurements were performed in a solution containing (v/v) 50% DMEM and 50% phosphate buffer (0.2 M, pH 7.4). The step potential was set as 20 mV, the pulse amplitude was at 50 mV, and the frequency was at 20 Hz. Based on the difference in the slope of the label-free TGT sensor and that of the labeled ones, the sensitivity difference was calculated to be ~140.

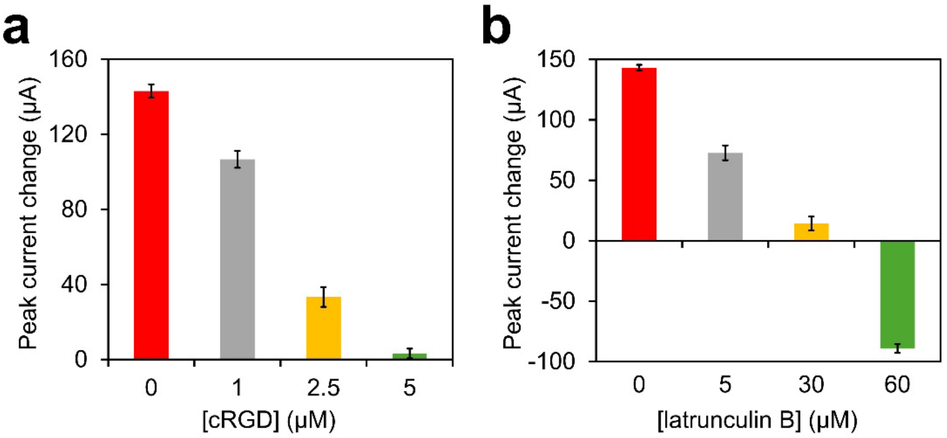


**Fig. S6.** **(a)** Peak current changes of the 12 pN TGT sensor after adding ~$1\times10^5$ HeLa cells for 75 min. These cells have been pre-treated with 0, 1, 2.5, or 5 μM of cyclic arginine-glycine-aspartic acid (cRGD) for 75 min. **(b)** Peak current changes of the 12 pN TGT sensor after adding ~$1\times10^5$ HeLa cells for 75 min. These cells have been pre-treated with 0, 5, 30, or 60 μM latrunculin B for 60 min. All these measurements were performed in a solution containing (v/v) 50% DMEM and 50% phosphate buffer (0.2 M, pH 7.4) and 5.0 mM $[Fe(CN)_6]^{4-}$. The step potential was set as 20 mV, the pulse amplitude was at 50 mV, and the frequency was at 20 Hz. Shown are the mean and standard error peak values after subtracting the background signals from four replicated tests.

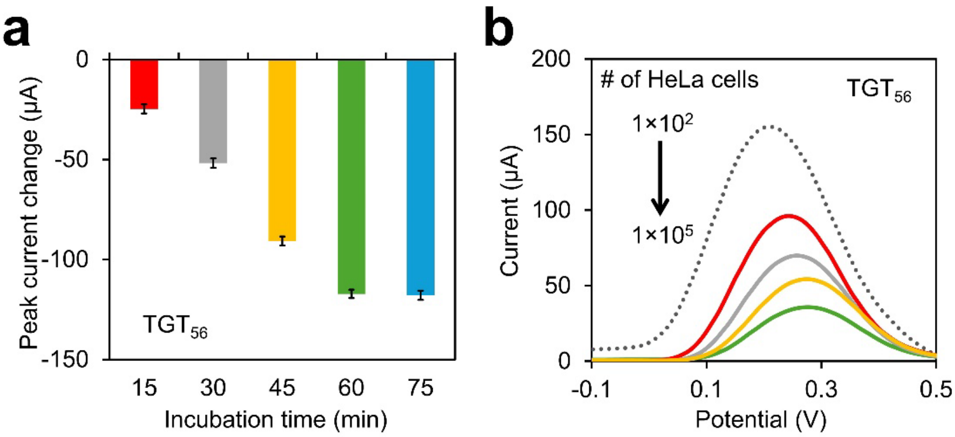


**Fig. S7.** **(a)** Peak current changes of the 56 pN TGT sensors after adding ~$1\times10^5$ HeLa cells for 15, 30, 45, and 60 min, respectively. **(b)** Square wave voltammetries of the 56 pN TGT sensors before (dotted line) and after adding ~100, ~$1\times10^3$, ~$1\times10^4$, or ~$1\times10^5$ HeLa cells for 60 min. All these measurements were performed in a solution containing (v/v) 50% DMEM and 50% phosphate buffer (0.2 M, pH 7.4) and 5.0 mM $[Fe(CN)_6]^{4-}$. The step potential was set as 20 mV, the pulse amplitude was at 50 mV, and the frequency was at 20 Hz. Shown are the mean and standard error peak values after subtracting the background signals from four replicated tests.

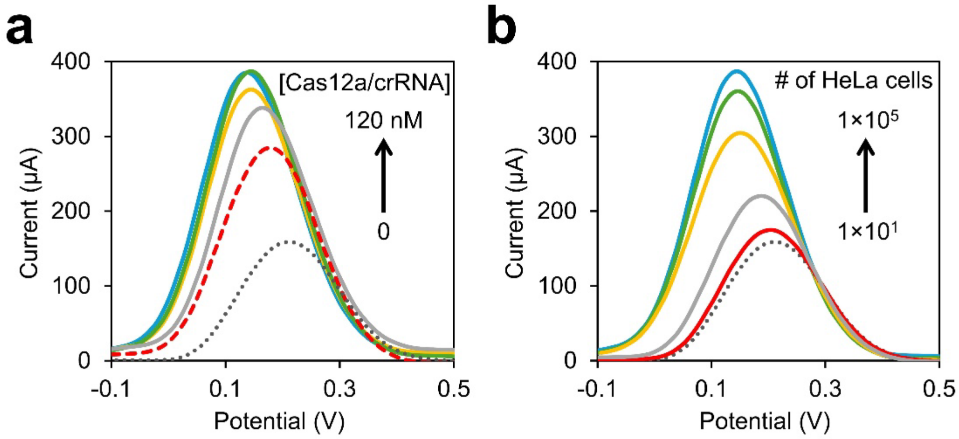


**Fig. S8.** **(a)** Square wave voltammetries of the 12 pN TGT sensors before (black dotted line) or after first incubating with ~$1\times10^5$ HeLa cells for 75 min, and then in the absence (red dash line) or adding 30, 60, 90, or 120 nM of the Cas12a/crRNA complex for 30 min. **(b)** Square wave voltammetries of the 12 pN TGT sensor before (black dotted line) or after adding ~10, ~100, ~$1\times10^3$, ~$1\times10^4$, or ~$1\times10^5$ HeLa cells for 75 min, in the presence of 90 nM Cas12a/crRNA complex. All these measurements were performed in a solution containing (v/v) 50% DMEM and 50% phosphate buffer (0.2 M, pH 7.4) and 5.0 mM $[Fe(CN)_6]^{4-}$. The step potential was set as 20 mV, the pulse amplitude was at 50 mV, and the frequency was at 20 Hz.

### Supplementary Table

**Table S1.** Parameters for determining the amount of doxorubicin in each TGT sensor.

| Molecules | Electron # | Q/Coulomb | $\Gamma$/mol·cm$^{-2}$ | Charge # | # per TGT |
|---|---|---|---|---|---|
| $[Ru(NH_3)_6]^{3+}$ | 1 | $4.05\times 10^{-6}$ | $3.5\times 10^{-10}$ | 3 | 42 |
| Doxorubicin | 2 | $1.85\times 10^{-6}$ | $8.0\times 10^{-11}$ | 1 | 3.2 |
